# Five decades of human H1N1 evolution reveal conserved glycan receptor specificity

**DOI:** 10.64898/2026.10.11.758004

**Authors:** Ruonan Liang, Jan-Willem Langenbach, Jadine Kuo, Andrei Bălas, Marju Zõbin, Geert-Jan Boons, Robert P. de Vries

**Affiliations:** Department of Chemical Biology & Drug Discovery, Utrecht Institute for Pharmaceutical Sciences, Utrecht University, Universiteitsweg 99, 3584CG, Utrecht, The Netherlands; Complex Carbohydrate Research Center, University of Georgia, 315 Riverbend Rd, Athens, GA 30602, USA

## Abstract

H1N1 influenza viruses have circulated continuously among human populations for five decades. During evolution, the hemagglutinin has accumulated amino acid substitutions in and around the receptor-binding site (RBS), mainly due to antigenic pressure, potentially altering receptor recognition and binding profiles. Here, we investigated the glycan-binding properties of H1N1 strains collected from 1977 through 2024 using ELISA, flow cytometry, and tissue histochemistry to characterize changes in receptor-binding specificity over time. Across the strains examined, we observe a recurring preference for binding α2,6-sialylated di-LacNAc N-glycans. Binding to α2,3-sialylated glycans was generally undetectable, although A/Hongkong/117/1977 and A/New Caledonia/20/1999 bind chicken tracheal tissues. Interestingly, post-pandemic H1N1 viruses fluctuate in their relative binding avidities. The responsible amino acids, in turn, have different effects in pre-pandemic HA proteins. We also report a classical egg-adapted H1N1 mutant that shows null binding across platforms. These findings highlight the contribution of RBS mutations during initial adaptation and evolution of human H1N1 influenza viruses to glycan-binding properties, which, in contrast to H3N2 viruses, never fail to bind glycans that correlate with avian erythrocyte hemagglutination.

## Introduction

Influenza A viruses have continuously circulated in humans for more than five decades. Among the three influenza A subtypes responsible for pandemics, H1N1 first caused the devastating 1918 influenza pandemic, resulting in high mortality worldwide. Seasonal H1N1 continued to circulate in humans until 1957, until the H2N2 subtype displaced it. H1N1 viruses, like those circulating in the 1950s, reemerged in 1977 via an accidental laboratory release^1^, and subsequently co-circulated with H3N2 viruses until 2009. In April 2009, a novel swine-origin influenza H1N1 virus, generated through triple reassortment from swine, humans, and birds^2,3^, became the dominant H1N1 lineage and was classified as pdmH1N1. Since 2009, all seasonal H1N1 viruses circulating in humans have descended from the pdmH1N1 lineage.

The hemagglutinin (HA) surface glycoprotein interacts with sialoside on the cell surface to infect the host through its receptor-binding site (RBS). The RBS, located on the globular head of HA, is composed of three domains that determine the specificity and affinity towards sialic acids: the 130-loop, the 190-helix, and the 220-loop^4^. In 1918, a human pandemic strain of HA carrying E190D and G225D switched binding specificity from avian-type α2-3 to human-type α2-6 sialylated receptors^5-7^. All post-1977 seasonal H1N1, pdm2009, and seasonal pdmH1N1 lineages with E190D/G225D maintained receptor-binding specificity to N-glycans, although early pdm2009 had low HA-receptor binding avidity^8^. Like H3N2 viruses, which showed a preference for binding to elongated glycans, H1 viruses have a strict preference for α2,6 sialylated glycans with di-LacNAc (Galβ1-4GlcNAc) structural motifs^9^. Recent research further confirmed that post-1977 and -pdm2009, and seasonal H1N1 viruses preferentially recognize α2,6 sialylated biantennary N-linked glycans containing two or more LacNAc repeats^10^.

H3N2 viruses have been shown to evolve to gain specific glycan-binding properties. During two periods, 2000-2019 and from 2025 onwards, these viruses fail to bind standard turkey erythrocytes^11-16^ used in vaccine development, hampering efficient antigenic analyses and vaccine selection. This specificity correlates with the lack of binding to omplex N-glycans, carrying di-LacNac repeats^11,13^. This phenotype has not been reported for H1N1 viruses, except for some egg-adapted strains. However, identifying H1 proteins with reduced receptor avidity is crucial for antigenic analyses.

Thus, defining which amino acids within or near the receptor-binding sites contribute to receptor-binding avidity in human H1N1 is crucial to potentially predict receptor-binding avidity. Here, we selected eight representative H1 strains spanning nearly five decades, from 1977 to the 2023/2024 season, to characterize their receptor-binding avidity. We then generated H1 proteins with amino acid substitutions at the receptor-binding sites to assess their effects on binding properties. Using tissue staining and flow cytometry (FACS), we evaluated their binding characterization biologically. We found that all selected strains bound α2,6 sialylated N-linked glycans with di-LacNAc antennae and bound human tracheal sections. Interestingly, A/Hongkong/117/1977 and A/New Caledonia/20/1999 can also bind chicken trachea. This is probably related to their expanded receptor-binding specificities. The egg-adapted strain A/Victoria/4879/2022 failed to bind receptors, but Q226R, present in the cell culture-derived vaccine strain WI/22, restored binding. During evolution, the effect of RBS residues on receptor-binding avidity depends strongly on genetic background.

## Results

### Changes in the binding properties of seasonal H1N1 during the pre-pandemic period (1977-2008)

Little is known about the receptor-binding characteristics of H1N1 viruses that emerged in the late 70s. To determine the binding avidity of human pre-pandemic H1 strains, α2,6-sialylated bi-antennary mono-(1), di-(2), and tri-LacNAc (3), and α2,3-sialylated bi-antennary tri-LacNAc (4) biotin-labeled N-glycans were tested by ELISA (Fig.1A). We selected a representative panel of recombinant HA proteins over time, consisting of A/Hongkong/117/1977(HK/77), a strain for when seasonal H1N1 was reintroduced in the human population in 1977, A/Memphis/7/1980(ME/80), a strain whose sequence identity was relative conserved from 1980 onwards, A/New Caledonia/20/1999(NC/99), which replaced previous H1N1 lineages and serves as a basis for seasonal influenza vaccines from 2000 onwards, and A/Kentucky/UR06-0538/2007(KY/07), representative of late New Caledonia-like H1N1 viruses circulating in North America before the 2009 pandemic. (Fig.1B). The HAs of all strains tested showed a strong binding avidity to α2,6-sialylated bi-antenna di-LacNAc (**2**) and tri-LacNAc (**3**) N-glycans. In addition, NC/99 and KY/07 showed strong binding to mono-LacNAc (**1**), whereas HK/77 and ME/80 showed almost no binding. No detectable binding to α2,3-sialylated glycan was observed (**4**), although NC/99 showed weak binding to α2,3-sialylated glycan (**4**) in the glycan array from previous research^9^.

Mardin-Darby Canine Kidney (MDCK) cells mainly display α2,3-sialylated structures with only a minor display of α2,6 on short N-glycans^17^. hCK cells mainly display α2,6-sialylated short N-glycans, whereas hCK-B3GNT2 cells display α2,6-sialylated on elongated N-glycans, respectively^18,19^. We characterized binding ability to these cell lines across all strains using flow cytometry (Fig. 1C). HK/77 bound all cell types tested. Compared with MDCK cells, HK/77 showed approximately 7-fold higher binding to hCK cells; furthermore, binding to hCK-B3GNT2 cells was around 2-fold higher than to hCK. In contrast, ME/80, as a descendant of HK/77, showed no detectable binding to MDCK cells, but retained strong binding to hCK, with binding to hCK-B3GNT2 cells approximately 2-fold higher than to hCK cells. NC/99 also bound all cell types. Compared with MDCK cells, NC/99 showed approximately 5-fold higher binding to hCK cells, while its binding to hCK-B3GNT2 cells was comparable to that of hCK cells. KY/07 strongly bound to hCK and hCK-B3GNT2 cells, with no detectable binding signal to MDCK cells. Overall, after H1N1 viruses were reintroduced before the 2009 pandemic, all strains tested showed a preference for binding to hCK cells, with stronger binding to hCK-B3GNT2 cells.

**Fig. 1.**
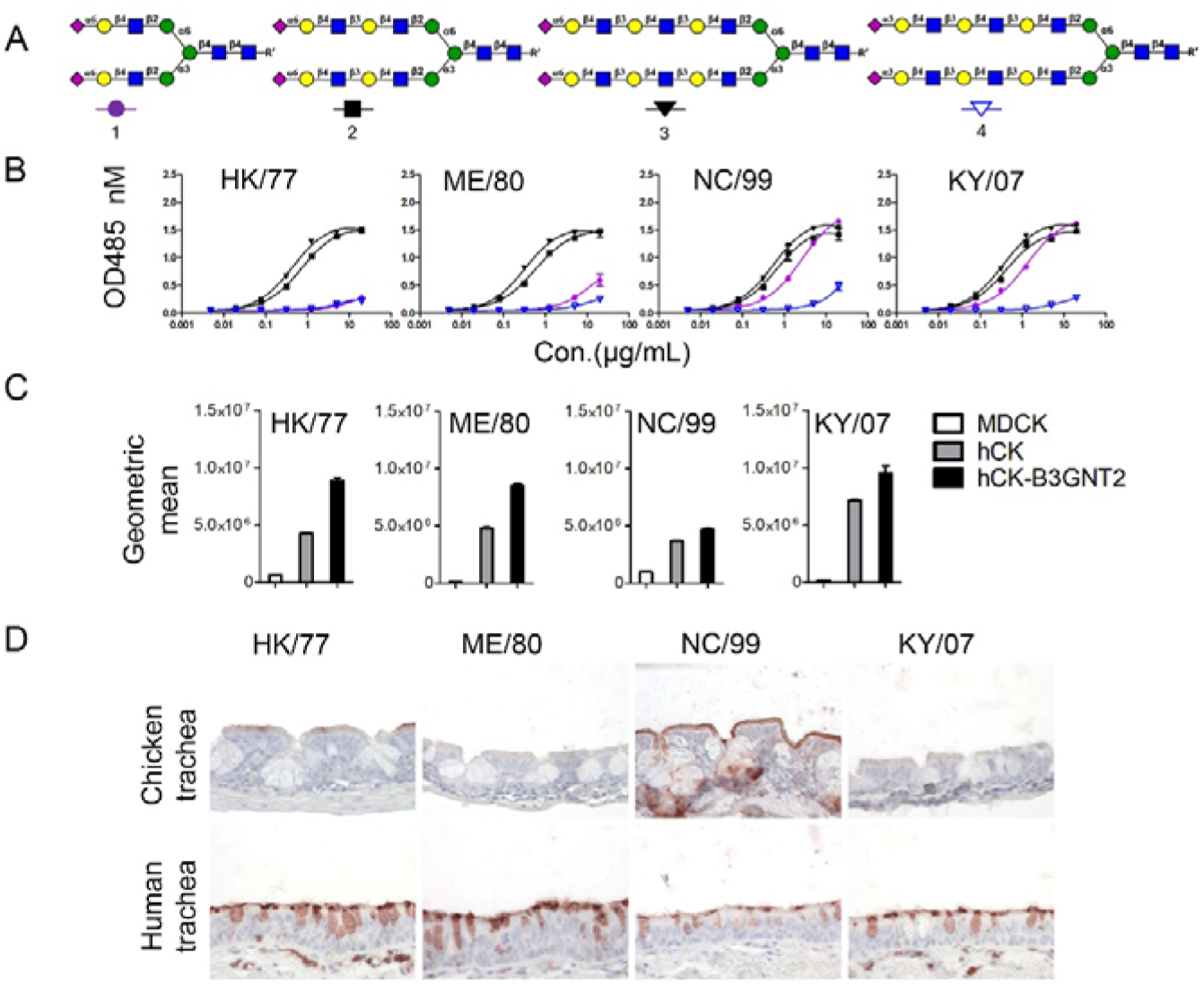
Change in the binding specificity of seasonal H1N1 during pre-pandemic period (1977-2008). A. N-Glycan profiles. B. Binding avidity of HK/77, ME/80, NC/99, and KY/07HAs to glycans was measured by ELISA. C. Binding of HK/77, ME/80, NC/99, and KY/07HAs proteins in MDCK, hCK and hCK-B3GNT2 cells were analyzed by flow cytometry. D. Binding of HK/77, ME/80, NC/99, and KY/07HAs to chicken and human tracheas were examined by tissue staining. For ELISA and flow cytometry assays, data were obtained from three independent experiments. Tissue staining experiments were performed in two independent experiments.

Chicken trachea tissue sections mainly display α2,3-sialylated N-glycans^20^, whereas human trachea is rich in α2–6 sialylated N-linked glycans with extended antennae^10^. In tissue histochemistry (Fig. 1D), HK/77 showed scattered binding signals on cilia of chicken tracheal epithelium, while NC/99 showed intense binding signals on cilia. In contrast, ME/80 and KY/07 showed no binding signal to chicken tracheal tissues. All strains showed binding signals on both cilia and goblet cells in the human trachea. In summary, all pre-pandemic H1N1 strains exhibited binding specificity to di- and tri-LacNAc with α2,6-linked sialic acid, hCK, and hCK-B3GNT2 cells. In addition, NC/99 and KY/07 bound the mono-LacNAc N-glycan, while HK/77 and NC/99 also showed detectable binding to MDCK cells and chicken tracheal tissues, indicating different receptor usage among the strains.

### E219K, G225D and A227E substitutions collectively alter HK/77 receptor binding specificity

It is well accepted that both D190 and D225 are vital for binding α2,6-linked sialic acid in H1 hemagglutinins, and that D225G is a likely egg adaptation^5,6,21^. We aligned the strains used in this study (Fig. S1), which shows that all strains carried D190 and shared several amino acid substitutions among ME/80, NC/99, and KY/07, including E219K, G225D, and A227E in receptor-binding sites. In addition, an amino acid deletion at position 133 appeared in the late 1990s. Here, we focus mainly on E219K, G225D, and A227E, as all three likely affect receptor binding and were quickly fixed in H1 after reintroduction into humans.

We introduced all single mutations in the HK/77 background, and ELISA results showed that HK/77 E219K exhibited binding avidity similar to HK/77, with preferential binding to di- and tri-LacNAc α2,6-sialylated compounds (**2** and **3**) (Fig. 2A). G225D retained binding to α2,6-SIA (**2** and **3**) and gained binding to the mono-LacNAc α2,6-sialylated glycan (**1)**. A227E retained binding to the tri-LacNAc (**3**) but showed decreased binding to the di-LacNAc α2,6-sialylated compound (**2**). This may be because E227 carries a bulky, negatively charged residue on the 220-loop^22^. None of the mutants showed detectable binding ability to α2,3-linked sialic acid on tri-LacNAc repeats (**4**). Surprisingly, flow cytometry and tissue histochemical staining (Fig. 2B and C) showed that E219K bound MDCK cells more strongly than HK/77, with binding to hCK and hCK-B3GNT2 cells nearly identical. HK/77 E219K bound cilia from chicken trachea and cilia and goblet cells from human trachea. We hypothesize that K219, a positively charged residue in the 220-loop, increases interaction with α2,6-sialylated structures. G225D, the hallmark amino acid, increases binding to MDCK, hCK, and hCK-B3GNT2 cells and binds only to human trachea. A227E significantly decreased overall binding avidity to hCK-B3GNT2 and failed to bind human trachea.

**Fig. 2.**
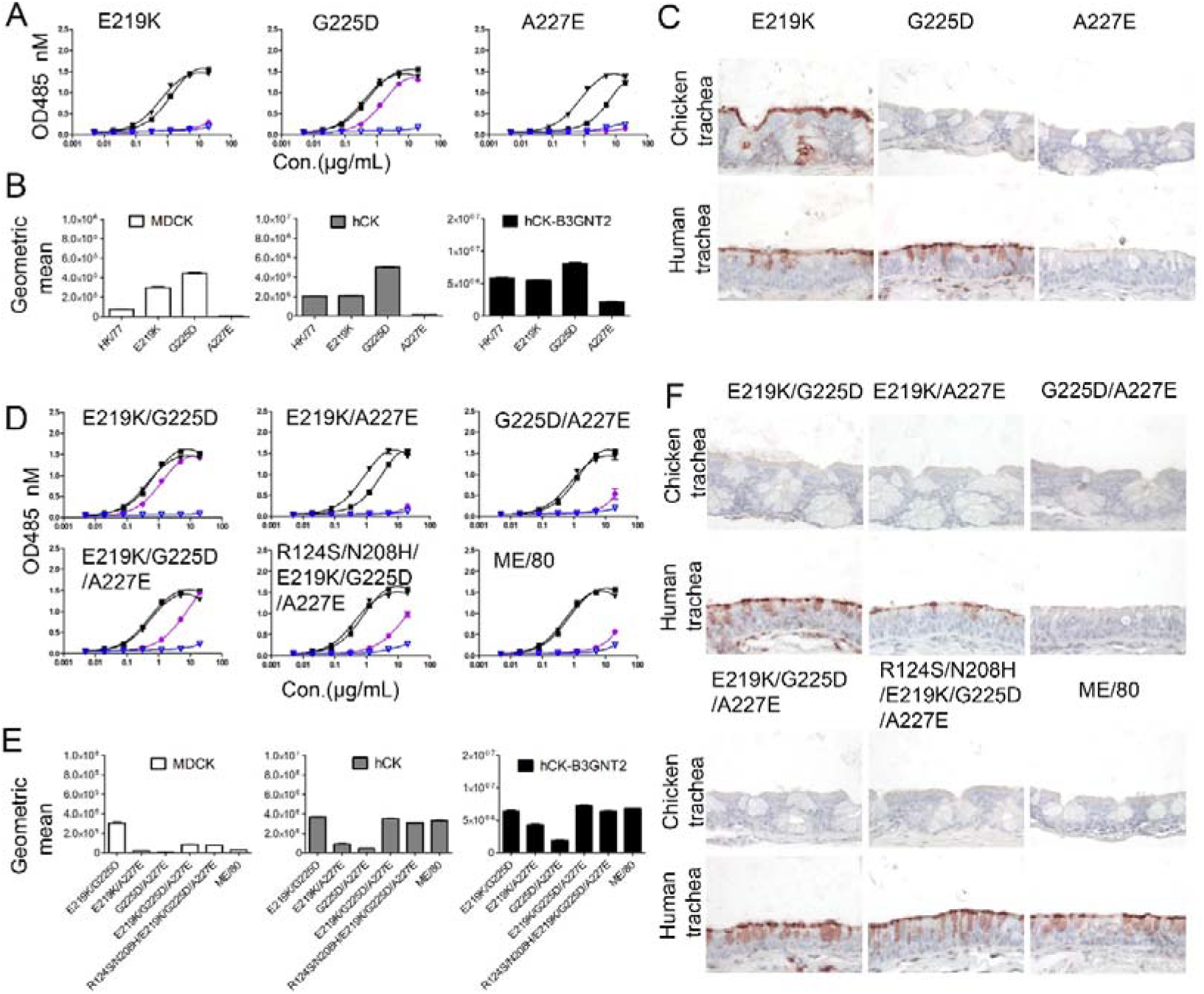
E219K, G225D, and A227E alter the receptor-binding profile of HK/77. A. and D. Binding avidity of HK77 mutants and ME/80 HAs to glycans was measured by ELISA. B. and E. Binding of HK/77 mutants and ME/80 in MDCK, hCK, and hCK-B3GNT2 cells was analyzed by flow cytometry. Binding levels were quantified based on the geometric mean fluorescence intensity. C and F. Binding of HK/77 mutants and ME/80 to chicken and human tracheas were examined by tissue staining. For ELISA and flow cytometry assays, data were obtained from three independent experiments. Tissue staining experiments were performed in two independent experiments.

Different combinations E219K/G225D, E219K/A227E, G225D/A227E, and triple mutation E219K/G225D/A227E were made to analyze the receptor binding properties. E219K/G225D, like G225D, bound all α2,6 sialylated N-glycans (**1, 2**, and **3**) (Fig. 2D). Adding A227E decreased overall receptor binding avidity in all backgrounds. E219K/A227E exhibited decreased binding to di-LacNAc (**2**), and G225D/A227E showed no detectable binding to α2,6 sialylated mono-LacNAc glycan (**1**) but retained binding to α2,6 sialylated di- and tri-LacNAc glycans (**2** and **3**). Interestingly, E219K/G225D/A227E regained binding activity to mono-LacNAc (**1**), although its binding appeared lower than that to di- and tri-LacNAc (**2** and **3**). ME/80 also carries R124S and N208H, which are near the RBS; we therefore generated HK/77 R124S/N208H/E219K/G225D/A227E. This mutant showed a similar binding profile for α2,6 sialylated di-LacNAc and tri-LacNAC glycans (**2** and **3**) but showed weaker binding to mono-LacNAc (**1**). Flow cytometry demonstrated that E219K/G225D, E219K/G225D/A227E, and R124S/N208H/E219K/G225D/A227E bound better to MDCK compared to the other HK/77 mutants, consistent with the ELISA results for binding the α2,6 sialylated mono-LacNAc glycan (**1)** (Fig. 2E). Whereas the HK/77 G225D/A227E and E219K/A227E mutants hardly bound to hCK and hCK-B3GNT2 cells, E219K/G225D, E219K/G225D/A227E, and R124S/N208H/E219K/G225D/A227E bound these cells efficiently. In histochemical staining of human and chicken tracheal sections, none of these mutants showed binding to chicken trachea (Fig. 2F). In contrast, they showed strong binding to the cilia and goblet cells of human trachea. G225D/A227E displayed a weaker signal, and E219K/A227E failed to bind either tissue.

Taken together, these results suggest that the combination of E219K, G225D, and A227E contributes to the receptor-binding profile of ME/80, with preferred binding to α2,6 sialylated di- and tri-LacNAc glycan and binding to human trachea tissues.

### Binding specificity of H1N1 after the 2009 pandemic

To evaluate the evolution of receptor-binding characteristics after the pandemic, the representative pdm09 lineage strains A/California/04/2009 (CA/04), A/Michigan/45/2015 (MI/15), A/Brisbane/02/2018 (BNE/18), and A/Wisconsin/67/2022 (WI/22) were tested. None of the strains showed detectable binding to the α2,3-sialylated structure (**4**). CA/04, as the earliest isolate, displayed weak binding to di- and tri-LacNAc glycans (**2** and **3**), as previously observed (Fig. 3A)^10^. We took CA/04 T200A/E227A as a control, as it has been reported to increase receptor-binding avidity compared with CA/04 ^8,23^. This protein exhibited increased binding to all three α2,6 sialylated glycans (**1, 2**, and **3**). MI/15 and BNE/18 showed strong binding to α2,6-sialylated di- and tri-LacNAc glycans (**2** and **3**). However, WI/22 displayed a stark decrease in binding to di- and tri-LacNAc glycans (**2** and **3**).

**Fig. 3.**
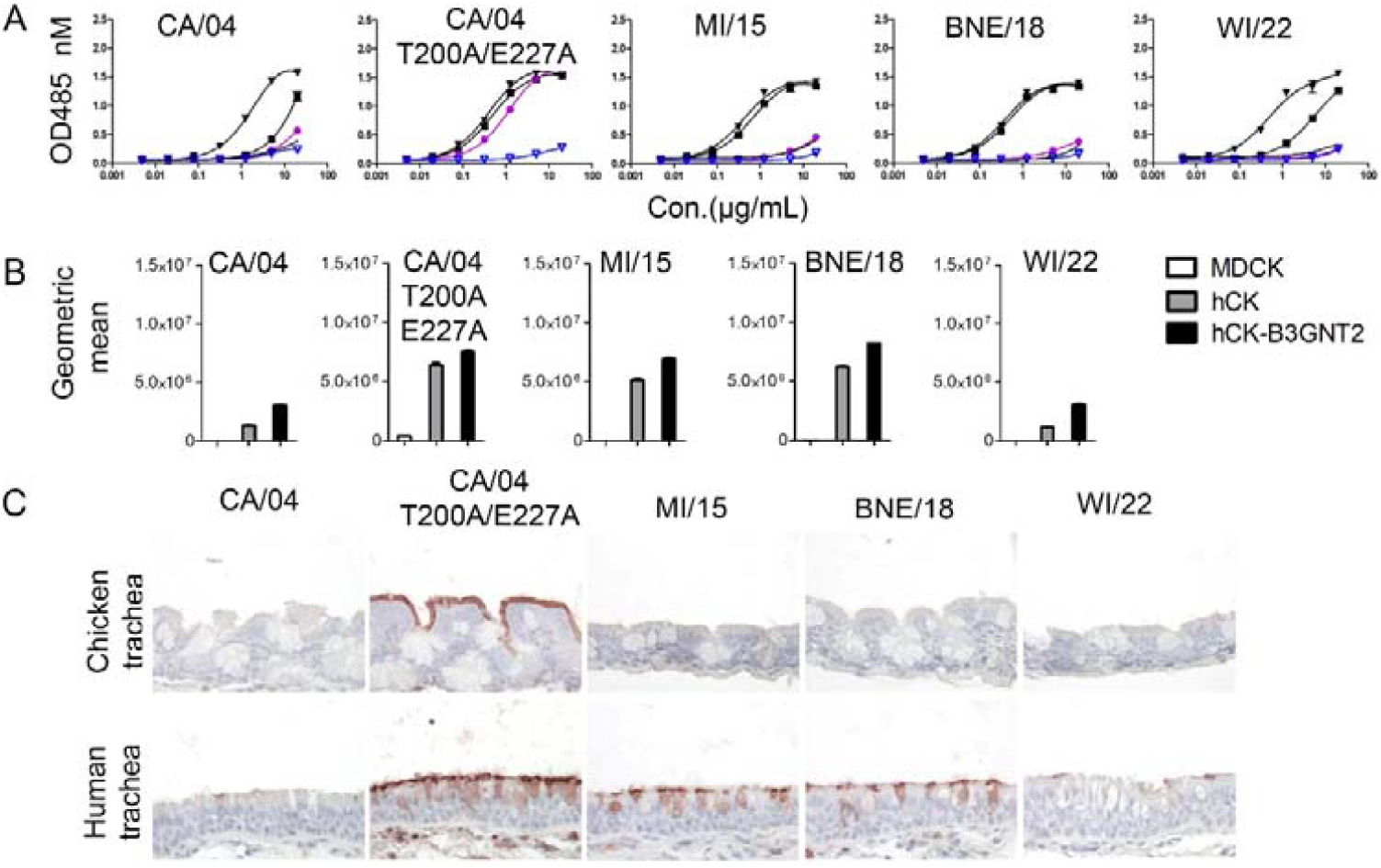
Binding specificity of seasonal H1N1 after the 2009 pandemic. A. Binding avidity of CA/04, CA/04 T200A/E227A, MI/15, BNE/18, and WI/22HAs to N-glycans was measured by ELISA. B. Binding properties of CA/04, CA/04 T200A/E227A, MI/15, BNE/18, and WI/22HAs proteins in MDCK, hCK and hCK-B3GNT2 cells were analyzed by flow cytometry. Binding levels were quantified based on the geometric mean fluorescence intensity. C. Binding properties of CA/04, CA/04 T200A/E227A, MI/15, BNE/18, and WI/22HAs to chicken trachea and human trachea were examined by tissue staining, For ELISA and flow cytometry assays, data were obtained from three independent experiments. Tissue staining experiments were performed in two independent

Flow cytometry showed that only CA/04 T200A/E227A bound to MDCK cells. All strains displayed different binding ability to MDCK, hCK, and hCK-B3GNT2 cells. CA/04 and WI/22 bound more weakly to both hCK and hCK-B3GNT2 cells, whereas CA/04 T200A/E227A, MI/15, and BNE/18 bound more strongly to hCK and hCK-B3GNT2. Notably, binding to hCK-B3GNT2 was stronger compared to binding to hCK cells (Fig. 3B). In histochemical staining, CA/04 T200A/E227A bound the cilia of chicken trachea and the cilia and goblet cells in human trachea (Fig. 3C), which may correspond to binding to compound **1**. In contrast, MI/15 and BNE/18 bound cilia and goblet cells in human trachea, CA/04 and WI/22 hardly bound human tracheal tissue. In summary, after the 2009 pandemic, the representative vaccine strains showed a preference for binding to α2,6-sialylated N-glycans with at least di-LacNAc motifs. However, WI/22 binds these compounds less avidly, affecting binding to hCK and hCK-B3GNT2 cells, as well as to human trachea.

### K133N and E227A alter glycan binding profiles in BNE/18

To investigate how amino acids substitution affect receptor binding properties between BNE/18 and WI/22, we performed a sequence alignment (Fig. S1) and identified four mutations, K133N, A189T, Q192E, and E227A, located within or near the receptor-binding site. In previous studies, D190, Q192 and S193 of A/South Carolina/1/1918 and A/Swine/Iowa/15/30 collectively interact with the GlcNAcβ1–3Galβ1–4Glcβ1-motif, of a sialylated ligand^21^. Additionally, T189 interacts with GlcNAc-5 through a water molecule in both the 1930-swine and 1934-human^24^. Amino acids at positions 190 and 193, as part of the 190-helex in H3N2 viruses, clearly affect receptor binding^25^. These structural observations prompted us to investigate whether the corresponding substitutions in BNE/18 affect glycan binding properties.

To determine the effects of K133N, A189T, Q192E, and E227A on glycan binding, we introduced these mutants individually into the BNE/18 background and analyzed the resulting proteins by ELISA. Unlike BNE/18 (Fig. 3A), K133N showed decreased binding avidity to α2,6-sialylated di- and tri-LacNAc glycans (**2** and **3**), whereas A189T and Q192E retained binding to both compounds (**2** and **3**). E227A showed binding to α2,6-sialylated mono-, di- and tri-LacNAc glycans (**1, 2** and **3**) (Fig. 4A). Using flow cytometry (Fig. 4B), we observed that among the protein tested, only the BNE/18 E227A protein bound to MDCK cells. The K133N mutant almost completely abolished binding to both hCK and hCK-B3GNT2 cells. In contrast, A189T, Q192E, and E227A bound to both cell types, although Q192E showed reduced binding compared to BNE/18 WT. In the tissue histochemistry, the BNE/18 E227A protein bound to chicken trachea, due to its ability to bind compound **1**. In contrast, BNE/18 K133N showed no detectable signal in either tissue. Both BNE/18 A189T and BNE/18 Q192E bound the human trachea, with the latter to a lesser extent (Fig. 4C).

**Fig. 4.**
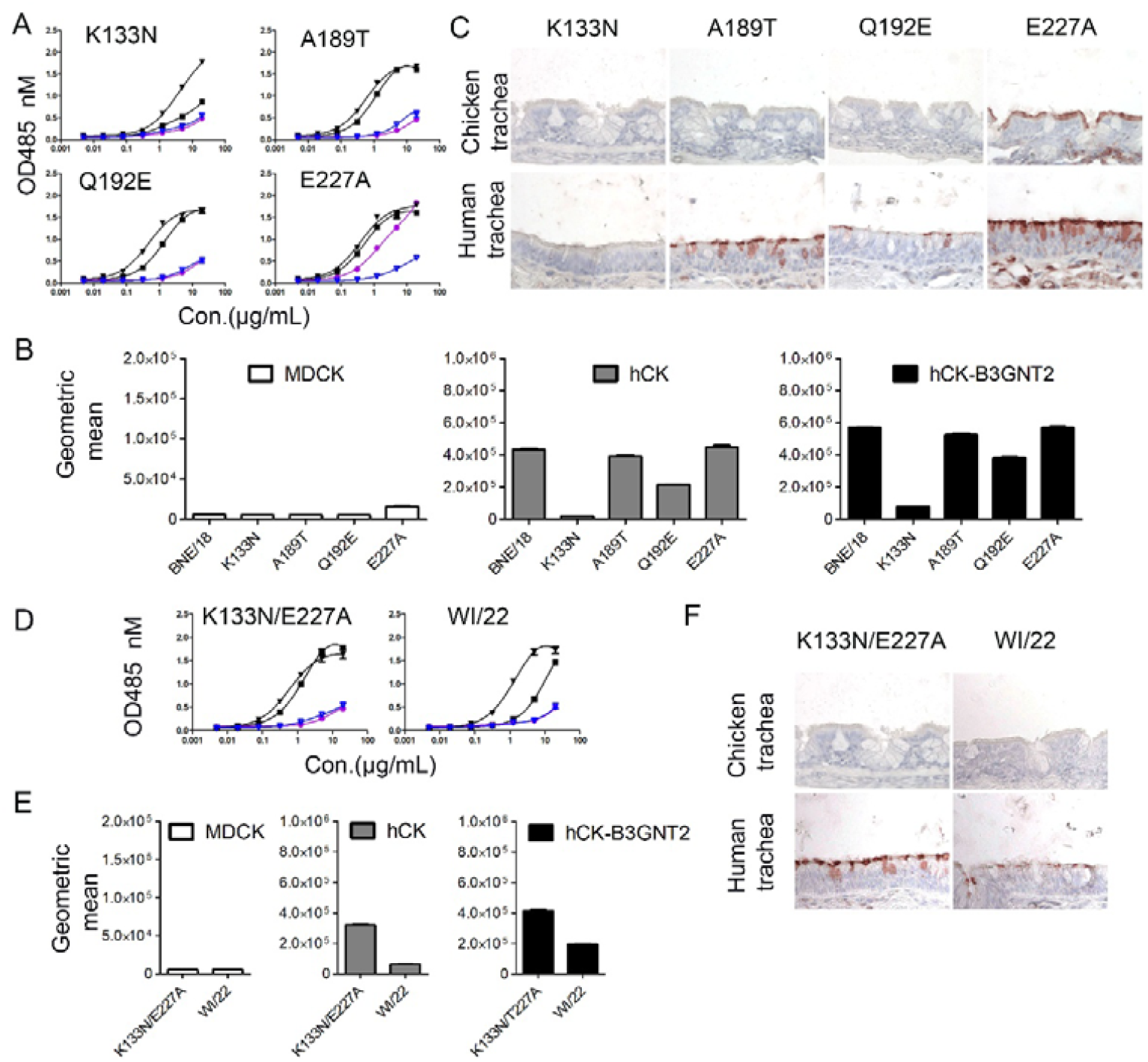
K133N and E227A alter glycan binding profiles in BNE/18. A. and D. Binding avidity of BNE/18 mutants and WI/22 to N-glycans was measured by ELISA. B. and E. Binding properties of BNE/18, BNE/18 mutants, and WI/22 HA proteins in MDCK, hCK, and hCK-B3GNT2 cells were analyzed by flow cytometry. Binding levels were quantified based on the geometric mean fluorescence intensity. C. and F. Binding properties of BNE/18 mutants and WI/22 to chicken and human tracheal tissues. For ELISA and flow cytometry assays, we obtained data from three independent experiments. We performed tissue staining experiments in two independent experiments.

Previously, amino acid deletion at 133 was reported in the late 1990s. We therefore removed R133 in HK/77 and ME/80. HK/77 R133Δ acquired binding to the 2,3-sialylated tri-LacNAc glycan (**4**) and showed an intense signal in chicken and human trachea. By contrast, ME/80 R133Δ bound 2,6-sialylated mono-, di-, and tri-LacNAc glycan (**1**) but did not gain binding to the chicken trachea (Fig. S2).

Because K133N and E227A significantly affected receptor binding properties, we generated a double mutant K133N/E227A. This mutant K133N/E227A bound only to 2,6-sialylated di- and tri-LacNAc compounds (**2** and **3**) (Fig. 4D), K133N/E227A exhibited binding ability to hCK and hCK-B3GNT2 cells, and an intense signal in the cilia and goblet cells of human trachea (Fig. 4E and 4F). These experiments show that K133N and E227A substitutions substantially alter receptor-binding avidity and specificity and contribute to binding to human tracheal tissues.

### Egg-adaptation in the 2025-26 NH H1N1 vaccine strain results in a loss of binding to biologically relevant substrates

A/Victoria/4897/2022 (VIC/22) and WI/22 were the recommended egg- and cell-adapted vaccines in the 2025-26 northern hemisphere influenza season, respectively. Unlike WI/22, we found that VIC/22 couldn’t bind α2,6-sialylated N-glycans (Fig. 5A). Correspondingly, this protein failed to bind MDCK, hCK, and hCK-B3GNT2 cells, and no detectable binding signal was observed in chicken and human trachea tissue sections (Fig. 5B and 5C). The A219T and R226Q substitutions differentiate the egg- and cell-adapted vaccine strains. We, thus, introduced A219T and R226Q into VIC/22 and observed that A219T did not increase receptor binding avidities of VIC/22(Fig. 5A); consistently, it failed to confer binding to MDCK, hCK, and hCK-B3GNT2 cells (Fig. 5B); and chicken trachea and human trachea (Fig. 5C). In contrast, R226Q rescued binding avidity to di- and tri-LacNAc compounds (**2** and **3**) (Fig. 5B); correspondingly, VIC/22 R226Q gained binding to hCK and hCK-B3GNT2 cells and displayed a binding signal on the cilia and goblet cells in human tracheal tissues (Fig. 5B and 5C), similarly to WI/22. Altogether, T219A and Q226R resulted in a loss of receptor binding during egg-culture adaptation.

**Fig. 5.**
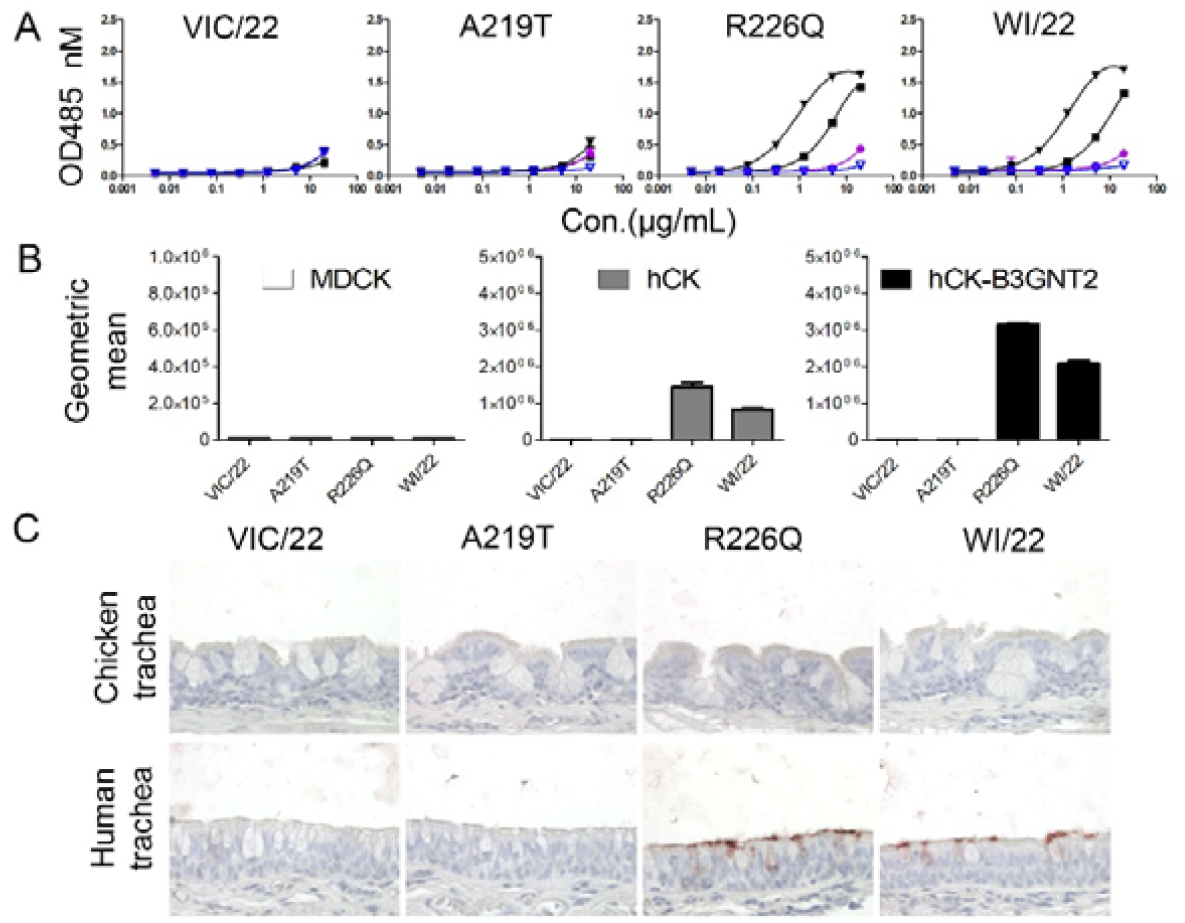
Egg-adaptation in the 2025-26 NH H1N1 vaccine strain results in a loss of binding. A. Binding avidity of VIC/22, mutants, and WI/22 to N-glycans was measured by ELISA. B. Binding properties of VIC/22, mutants, and WI/22 in MDCK, hCK, and hCK-B3GNT2 cells were analyzed by flow cytometry. Binding levels were quantified as geometric mean fluorescence intensity. C. Binding properties of VIC/22, mutants, and WI/22 to chicken and human trachea were examined by tissue staining. For ELISA and flow cytometry assays, we obtained data from three independent experiments with N=2 or N=3 replicates. We performed tissue staining experiments in two independent experiments.

## Discussion

In this study, we analyzed the receptor-binding specificity of human H1N1 influenza viruses from 1977 and 2024, all of which prefer α2,6 sialylated glycans with at least a di-LacNAc motif. In the late 1990s and before 2009, seasonal strains NC/99 and KY/07 bound mono-LacNAc and showed different binding properties in cells and tissues. Consistent with previous research, NC/99 binding to chicken trachea might depend on binding to simple N-glycans with α2,6-linked sialic acids. However, KY/07 doesn’t bind to chicken trachea, although it efficiently binds to α2,6-sialylated mono-LacNAc glycans. In contrast with recent literature^10^, we do not observe any α2,3-linked sialic acid binding by NC/99, which may be attributed to experimental details. Thus, low α2,3-linked sialic acid binding by NC/99 may explain its efficient binding to chicken tracheal tissues^9^.

Amino acid 133, which is either mutated or deleted in most H1N1 viruses, has been identified as an important residue for recognition by human monoclonal antibodies^26,27^. In our study, K133N in BNE/18 decreased receptor-binding avidity, consistent with the effect reported for A/California/07/2009 K133N^27^. Deleting amino acid 133 in HK/77 and ME/80 produced different receptor-binding profiles; these phenotypes suggest that alterations at residue 133 can affect receptor-binding avidity and specificity in a genetic-background-dependent manner, similar to the effect we previously observed for H5N1 receptor binding^28,29^. Residues 131, 132, and 133 form a structural cluster in the receptor-binding pocket^30^, suggesting that substitutions at 133 may change the structural conformation and consequently affect binding.

Influenza antigenic drift can affect receptor-binding avidity^31^, which will likely depend on the genetic backbone. We foresee a similar differential effect when positions 219, 225, 226, and 227 are mutated in different backgrounds. Indeed, position 227 changes from A to E and vice versa at different points during evolution, likely balancing receptor-binding avidity and HA-NA balance^32^. In CA/04, E227A together with T200A increased receptor binding to α2,6 sialylated glycans, whereas A227E in HK/77 reduced receptor binding to α2,6 sialylated di-LacNAc compound and exhibited additional effects in the presence of G225D/E219K. Similarly, E227A in BNE/18 works with K133N to modulate receptor binding. These results suggest that residue 227 may not function independently but instead interact with other HA residues to regulate receptor binding avidity. The different effects of residue 227 in these genetic backgrounds further suggest that receptor-binding properties depend on a combination of multiple HA residues rather than a single substitution. This is consistent with previous studies that the effects of substitutions can depend on the surrounding HA sequence context^33^.

Indeed, in egg-grown 2009 pandemic H1N1 viruses, Q226R is one of the most frequent changes in the receptor-binding pocket and is known to impact receptor binding^34^. Previous studies showed that it can change receptor binding preference from human-type α2,6 sialylated glycans toward avian-type α2,3 sialylated glycans^35-37^. In our study, Vic/22, which contains R226 and D225, completely abolishes binding to α2,6-sialylated glycans and fails to bind to human and chicken trachea. D225 may explain this phenotype, as it can influence the conformation of the 220-loop and thereby affect the position and receptor interaction of residue 226^38^.

Epistatic effects of amino acids in and around the RBS are an accepted phenomenon in H3N2 viruses^39-41^, but less so in H1N1. Here we report several examples of amino acid mutations that arise under antigenic pressure and steer receptor binding toward different structures depending on the genetic background.

## Materials and methods

### HA expression and purification

HA encoding cDNAs of A/Hongkong/117/1977, A/Memphis/7/1980, A/New Caledonia/20/1999, A/Kentucky/UR06-0538/2007, A/California/04/2009, A/Michigan/45/2015, A/Brisbane/02/2018, and A/Victoria/4897/2022 were cloned into the pCD5 expression vector, with a signal sequence, GCN4 trimerization motif((KQIEDKIEEIESKQKKIENEIARIKK), a TEV cleavage site, the twin-strep (WSHPQFEKGGGSGGGSWSHPQFEK, IBA, Germany) and a mOrange2. Mutations in HAs were generated using site-directed mutagenesis. The HAs were expressed in the HEK 293S GnTI(-) and purified from cell culture supernatants as described earlier^42^. In brief, HEK 293S GnTI(-) was transfected using the expression vectors with polyethyleneimine I (PEI) (DNA:PEI, 1:8). 6 hours post-transfection, the transfection medium was removed and replaced by the 293 SFM II expression medium (Gibco), supplemented with 3.0 g/L Primatone (Kerry), 2.0 g/L glucose, 3.6 g/L bicarbonate, 0.4 g/L valproic acid, 1% glutaMAX (Gibco), and 1.5% DMSO. After -5days of incubation, supernatants were collected. HAs were purified using Strep-Tactin®XT 4Flow® resin (IBA, Germany) according to the manufacturer’s instructions.

### Enzyme-linked immunosorbent assay (ELISA)

Nunc Maxisorp 96-well plates (Invitrogen) were coated with streptavidin (50 μL, 5 μg/mL, Westburg) at 4°C in PBS overnight, followed by blocking with 300 μL of 1% BSA in PBS-T for 3 hours at room temperature. The streptavidin-coated plates were coated with 50 μL of 50 nM biotinylated glycans overnight at 4°C, followed by blocking with 1% BSA in PBS with 0.1% Tween-20 (PBS-T). HAs (20 μg/ml were precomplexed with strep-mab (10 μg/ml) and goat-anti-human antibodies (5 μg/ml) Invitrogen, #31410) to afford a 1:0.65:0.325 molar ratio; this mix is then incubated on ice for 30 min. Pre-complexed proteins were added to the first row (75 μL), diluted serially 1:3, and incubated for 90 min at room temperature. HA binding was developed with OPD and stopped with 2.5 M H_2_SO_4_ (100 μL) after 5 min. Optical density was recorded at 485 nm (Polar Star Omega, BMG Labtech). Experiments were performed independently three times, with each experiment in duplicate.

### Protein histochemistry

Protein histochemistry was performed as previously described^43^. Briefly, 5 μm chicken and human tracheal sections were deparaffinized and rehydrated. The endogenous peroxidase was inactivated by 1% H_2_O_2_ in MeOH for 30 min at room temperature. Slides were washed with PBS-T. Tissues were blocked with 3% BSA in PBS overnight at 4°C. The precomplexed HA, strep-mab, and goat-anti-human antibodies were incubated with tissues for 90 min at room temperature. HA was 10 ug/ml. After incubation, the tissue was stained with AEC substrate for HRP (Abcam) and subsequently stained with hematoxylin.

### Flow cytometry analysis

10 μg/mL Hemagglutinin was precomplexed with strep-mab and goat anti-human Alexa Fluor 488 at a 1:0.65:0.325 molar ratio on ice for 30 min. The precomplexed proteins were incubated with MDCK, hCK, or hCK-B3GNT2 cells for 30 min at 4°C in the dark. Cells were washed with FACS buffer (PBS containing 1% FCS and 2 mM EDTA) and followed by the centrifuge at 300 rcf for 5 min. The cells were fixed with 1% Paraformaldehyde for 10 min at 4°C in the dark. After being washed with FACS buffer, the cells were resuspended in 100 μL of FACS buffer. Flow cytometry was performed using the CytoFLEX LX (Beckman Coulter). Data were analyzed using FlowJo software. All cells, single cells, and live cells were gated. Mean fluorescence values for triplicates were averaged, and standard deviations were calculated.

## Acknowledgements

R.L. is supported by a CSC fellowship (202209120001).

## Supplementary

**Fig. S1.**
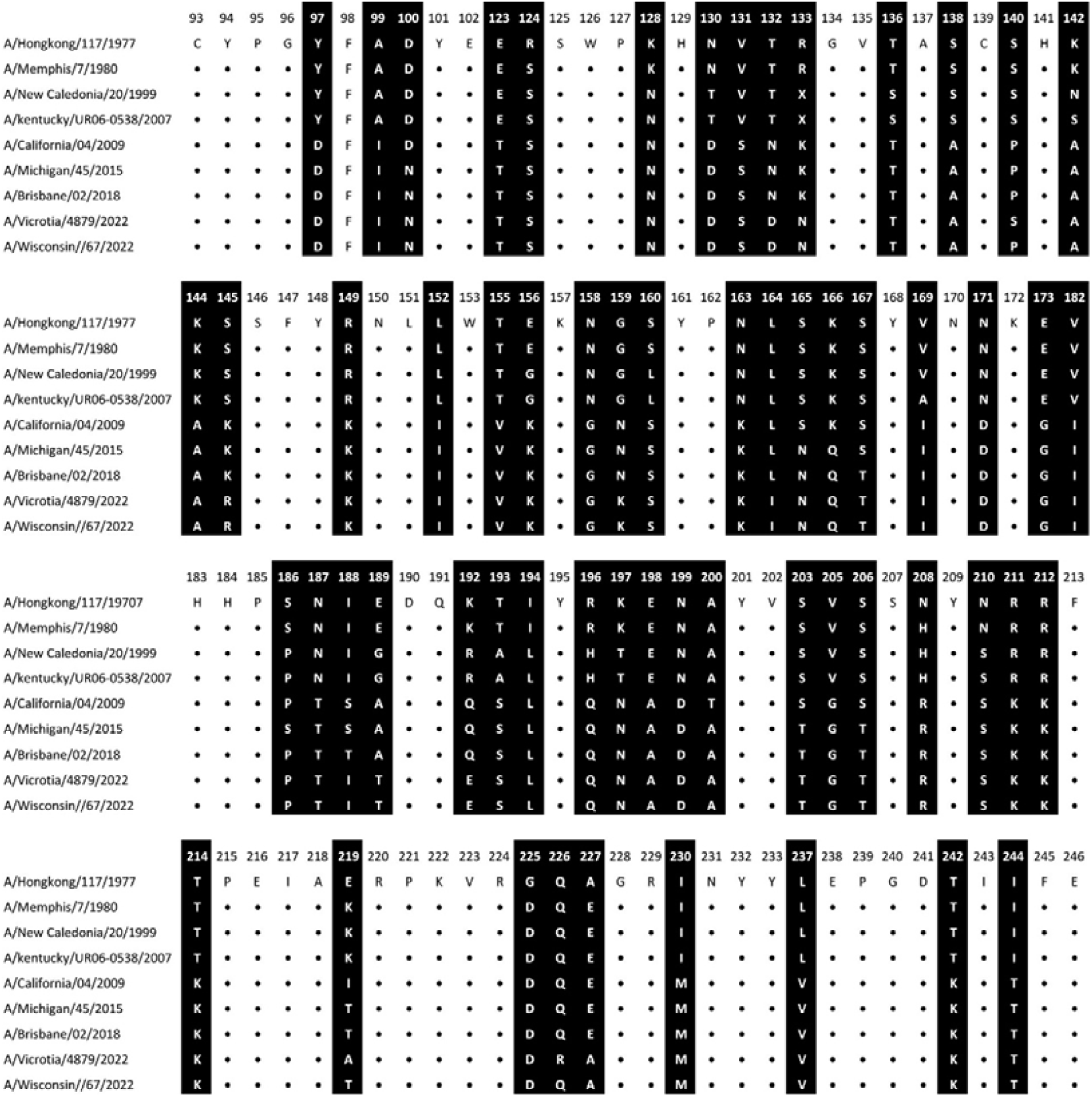
Sequence alignment of theH1N1 HA sequence spanning from 1977 to 2023/2024 season.

**Fig. S2.**
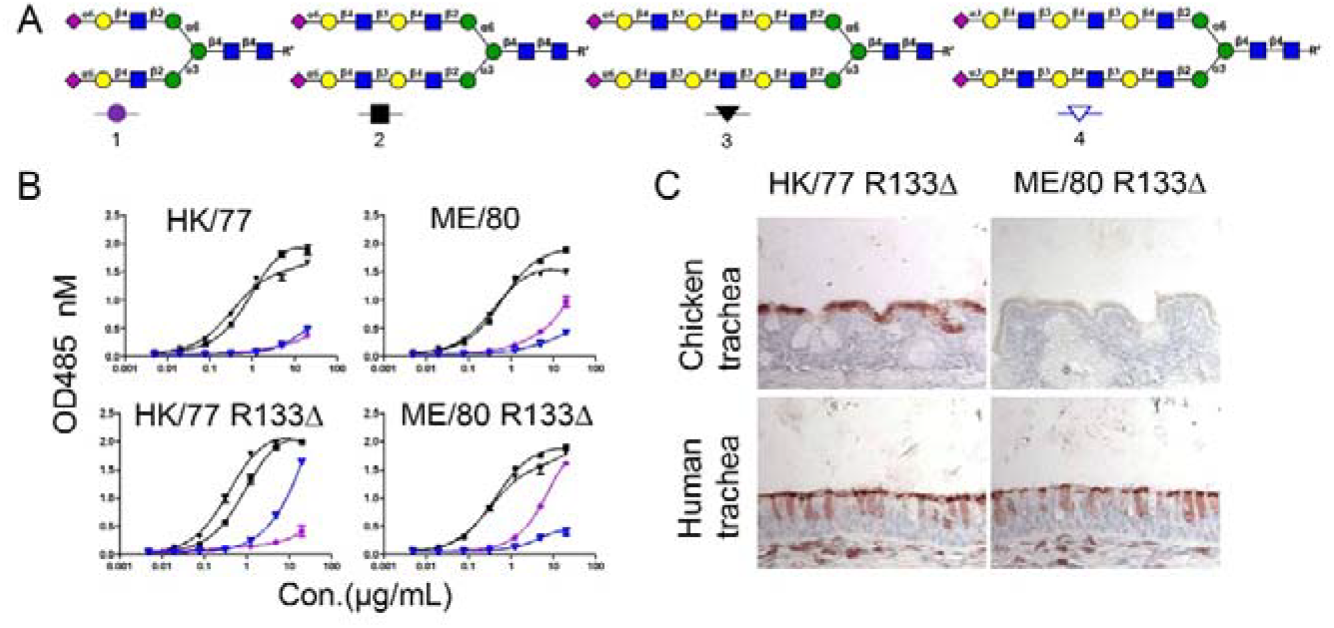
Binding ability of deletion of 133 in HK/77 and ME/80. A. N-Glycan profile. B. Binding avidity of HK/77 WT, K133_Δ_, and ME/80 WT, K133_Δ_ to N-glycans was measured by ELISA. C. Binding properties of HK/77 WT, K133_Δ_, and ME/80 WT, K133_Δ_ to chicken and human trachea were examined by tissue staining. For ELISA assays, data were obtained from three independent experiments. Tissue staining experiments were performed in two independent experiments.

